# Within and among population differences in cuticular hydrocarbons in the seabird tick *Ixodes uriae*

**DOI:** 10.1101/2022.01.21.477272

**Authors:** Marlène Dupraz, Chloe Leroy, Thorkell Lindberg Thórarinsson, Patrizia d’Ettorre, Karen D. McCoy

## Abstract

The hydrophobic layer of the arthropod cuticle acts to maintain water balance, but can also serve to transmit chemical signals via cuticular hydrocarbons (CHC), essential mediators of arthropod behavior. CHC signatures typically vary qualitatively among species, but also quantitatively among populations within a species, and have been used as taxonomic tools to differentiate species or populations in a variety of taxa. Most work in this area to date has focused on insects, with little known for other arthropod groups such as ticks. The worldwide distribution and extensive host-range of the seabird tick *Ixodes uriae* make it a good model to study the factors influencing CHC composition. Genetically differentiated host-races of *I. uriae* have evolved across the distribution of this species but the factors promoting sympatric population divergence are still unknown. To test for a potential role of host-associated CHC in population isolation, we collected *I. uriae* specimens from two of its seabird hosts, the Atlantic puffin (*Fratercula arctica)* and the common guillemot (*Uria aalge*) in different colonies in Iceland. Using gas-chromatography and mass-spectrometry, we detected a complex cuticular mixture of 22 hydrocarbons, including *n*-alkanes, methyl-alkanes and alkenes ranging from 17 to 33 carbons in length. We found that each population had a distinct CHC profile. The host group explained the greatest amount of population divergence, with long-chain hydrocarbons being more abundant in puffin tick populations compared to guillemot tick populations. Future work will now be required to test whether the different CHC signals reinforce assortative mating, thereby playing a role in generating *I. uriae* population divergence patterns, and to evaluate diverse hypotheses on the origin of distinct population signatures.

## Introduction

The arthropod cuticle acts as both an exoskeleton and a barrier from the external environment (Andersen, 1979). The outermost layer, the epicuticle, is covered by a lipid layer (Lockey, 1988) made up of esters, carboxylic acids, alcohols, carbonyls and long-chain hydrocarbons (Andersen, 1979) that protects the organism from desiccation (Filshie, 1982). However, cuticular hydrocarbons (CHC) are also involved in chemical communication, serving as sex pheromones, kairomones and/or signature mixtures allowing recognition of social identity (van Zweden and d’Ettorre, 2010; Wyatt, 2010). Using analytic techniques such as gas chromatography-mass spectrometry (GC-MS) and MALDI-TOF mass spectrometry, researchers have described hydrocarbons of up to 70 carbons in chain length in insects, principally *n*-alkanes, methyl-branched alkanes and alkenes (Blomquist and Bagnères, 2010). The array of CHC on the cuticle constitute a species-specific chemical signature, varying qualitatively between species and quantitatively within species (Lockey, 1988). CHC patterns are genetically controlled, but the relative abundance of particular components can be linked to environmental conditions (Estrada-Peña et al., 1993; Gibbs et al., 1991). CHC patterns have been used as taxonomic tools to characterize hundreds of arthropod species (Howard and Blomquist, 2005), and to discriminate closely-related populations (Bagnères et al., 1991; Bartelt et al., 1986; Jallon and David, 1987; Kruger et al., 1991; Simmons et al., 2014).

Ticks are hematophagous arthropods widely distributed across the globe and parasitize a diverse array of vertebrate species (McCoy and Boulanger, 2015). However, little work has been performed to describe CHC profiles and their variation among species and populations.The only studies to date have used CHC profiles in an attempt to differentiate closely-related taxa (Estrada-Peña et al., 1992, 1994, 1996; Hunt, 1986; Estrada-Peña and Dusbabek, 1993). However, tick CHCs may play essential roles in several aspects of tick life histories. First, the tick epicuticle is perforated with numerous channels providing a large surface for exchange with the external environment. As most tick species spend the major part of their life cycle in the off-host environment, maintaining water balance across this surface under different environmental conditions will directly dictate survival (Randolph and Storey, 1999); the presence of CHCs likely plays an important role in this. Second, as obligate parasites, access to the vertebrate host for the bloodmeal is a key aspect of the tick life cycle and, as such, ticks have adapted key traits to locate and successfully exploit their host (McCoy and Boulanger, 2015), as for example Haller’s organ, a unique chemosensory structure located on the 1st pairs of legs (Leonovich, 2004). Shimshoni et al. (2013) found *Rhipicephalus* tick species had different cuticular fatty acid compositions in relation to host use. However, whether these differences are linked to adaptive survival or are by-product variation due to the host resource is unknown as yet. Finally, ticks aggregate both on hosts and in the off-host environment (Randolph, 1998). This behavior is thought to facilitate blood feeding on the host and increase survival in the off-host environment. It may also enable ticks to find appropriate mates for reproduction. Some ticks, such as the tropical bont tick *Amblyomma variegatum*, produce a multicomponent pheromone to provoke aggregation (Schöni et al., 1984) but the potential role of these pheromones in assortative mating has never been examined.

Seabird ticks are generally nidicolous, exploiting different local host species that use diverse micro-habitats within the colony (Dietrich et al., 2011). Due to this diversity, these ticks may experience diverse selective pressures coming from both the hosts and from the temperature and humidity conditions of the nest micro-habitat. In particular, *Ixodes uriae*, a tick associated with seabird colonies in the polar regions of both hemispheres, is known to form host-specific races that show genetic (McCoy et al., 2005, 2003, 2001) and morphological (Dietrich et al., 2013) differences in relation to host use. Differential performance on alternative hosts has also been experimentally demonstrated (Dietrich et al., 2014). Nevertheless, the factors driving divergence within this species have yet to be specifically identified. A potential role for isolating mechanisms that lead to assortative mating based on host use have been suggested (McCoy et al., 2013).

Here, we examine the degree of cuticular hydrocarbon diversity in *I. uriae* and the factors influencing these chemical signatures. Based on current knowledge, we predicted that cuticular hydrocarbon patterns could vary in ticks 1) exploiting different host species and 2) sampled in different geographic locations. If host exploitation modifies CHC diversity and abundance, we expected that signatures in ticks from the same seabird host in different locations should be more similar than signatures in ticks from different host species in the same geographic location. To test these predictions, we collected *I. uriae* specimens in the nesting area of two seabird host species, the common guillemot (CG) *Uria aalge*, and the Atlantic puffin (PF) *Fratercula arctica*, at three locations in Iceland. We then extracted cuticular hydrocarbons and analyzed them using gas chromatography-mass spectrometry (GC-MS).

## Methods

### Population samples

Flat adult female ticks were collected off-host in three sites in Iceland in June 2016: common guillemot ticks (CG) were collected under rocks in the middle of the colony in Langanes (L) (66°22′07.2″N 14°38′33.0″W) and on Grimsey Island (GRI) (66°32′57.3″N 17°59′31.1″W); puffin ticks (PF) were collected in burrows on Lundey Island (LI) (66°06′53.2″N 17°22′13.1″W) and on Grimsey Island (GRI) (66°32′39.1″N 18°01′12.9″W) (Fig.1). Ticks were kept alive in plastic tubes before hydrocarbon extraction. Four replicates of 10 flat ticks were extracted for each host species at each site.

**Figure 1:**
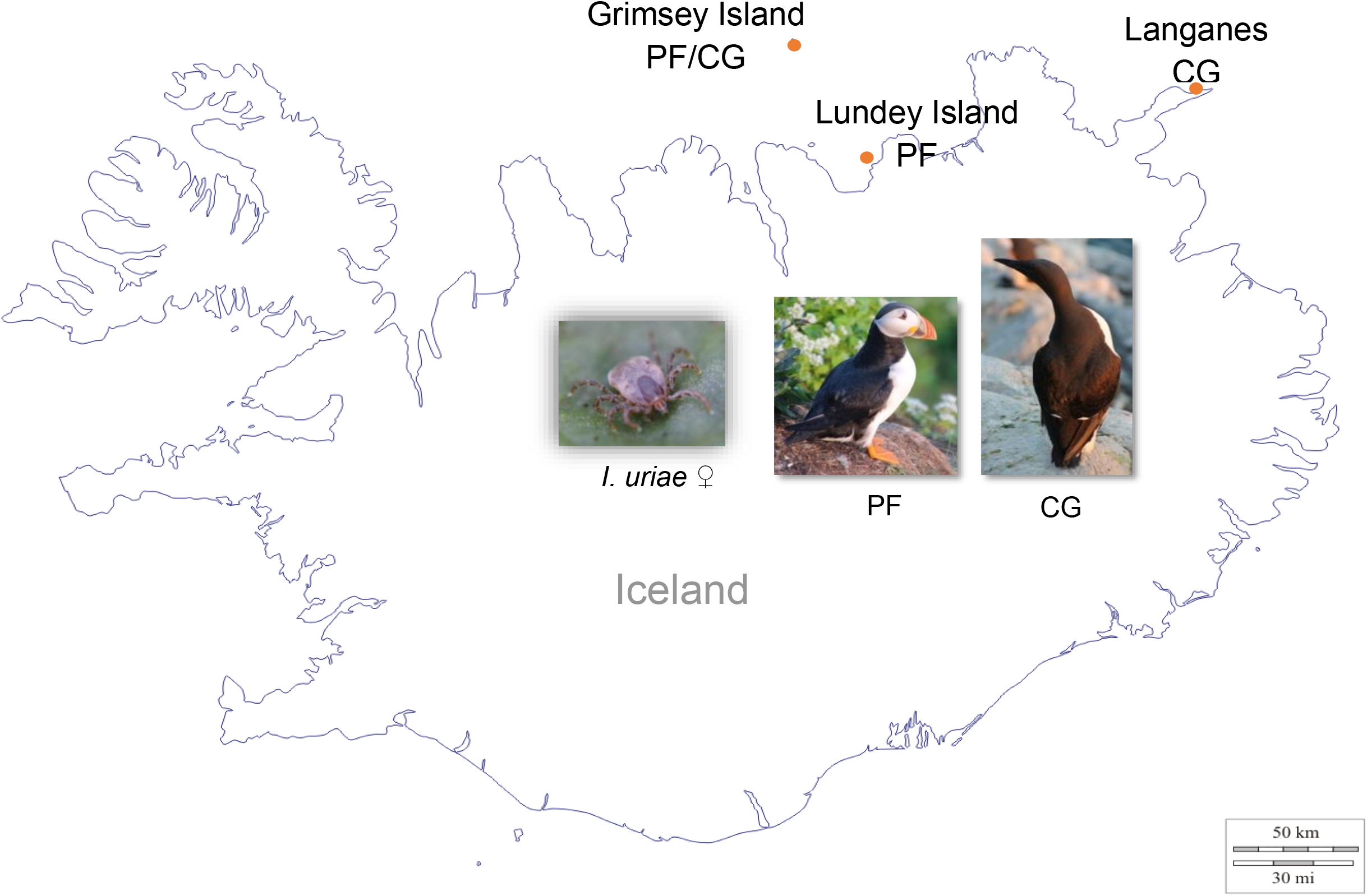
Map showing the sampling locations for *Ixodes uriae* ticks and hosts in Iceland in June 2016. PF refers to the Atlantic puffin (*Fratercula arctica*) and CG to the common guillemot (*Uria aalge*). The acronyms of the sampling locations are as follows: Grimsey Island - G; Lundey Island – Li; Langanes - L.

### Hydrocarbon extraction

Several extraction attempts were made with ticks kept alive or stored in alcohol and with only one individual or in batches of five, but the quantities of extracted cuticular compounds were too low to use in analyses. Cuticular compounds were therefore extracted by immersing 10 living flat female ticks in 200 μl of pentane (HPLC grade, Sigma-Aldrich) in glass vials. The vials were agitated for 1 minute, set down to rest for 5 minutes and then re-agitated for 1 minute. Ticks were then removed from the vials and preserved in 70% ethanol. The pentane was then evaporated off over 30 minutes to obtain dried eluent and the vials were closed and kept at 4°C until analysis. A negative control containing only 200 μl of pentane was added to each extraction session to control for potential contamination. We did not use an internal standard because the aim of our analysis was not to quantify the absolute amount of CHCs, but to compare the relative proportions of CHCs among samples.

### Chemical analyses

Cuticular hydrocarbons are not very volatile, but to reduce variation during the evaporation process, all samples were treated in the same way, placed under the same fume hood in a temperature controlled room. Samples were re-diluted in 40μl of pentane and 3μl were injected into an Agilent Technologies 7890A gas chromatograph (capillary column: Agilent HP-5MS, 30 m × 0.25 mm × 0.25 μm; split–splitless injector; carrying helium gas at 1 mL/min) coupled with an Agilent 5975C mass spectrometer with 70 eV electron impact ionization. The oven temperature was programmed at 70°C for 1min, and was increased at 30°C/min to 200°C, then to 320°C at 5°C/min and held for 5 min. Compounds were then identified (raw data available on Zenodo at https://doi.org/10.5281/zenodo.6497483) and sorted on the basis of their mass spectra and retention time by comparison with standards (alkane standard solutions, Sigma-Aldrich, Saint Louis, MO, USA) and published spectra (NIST Library). The abundance of each detected cuticular hydrocarbon was used to build a box plot using the ggplot 2 package with R software (version 4.1.1)(script is available on Zenodo at https://doi.org/10.5281/zenodo.7018260). The areas under the peaks were extracted using the Agilent MSD ChemStation software (E.02.01.1177) and were used to calculate the relative amount of each hydrocarbon to normalize data coming from different origins and of different qualities. The mean peak area of the four population replicates was calculated for each hydrocarbon to obtain a colony average.

### Multivariate analyses

For analytical purposes, if the relative abundance of a CHC was 0, the value was replaced by 0.00001 which is several times lower than the lowest quantity found for a CHC in our dataset. The data were then transformed using a centered log ratio procedure (data available in supplementary materials at https://doi.org/10.5281/zenodo.5889077) and analyzed by Partial Least Squares coupled with a Discriminant analysis (PLS-DA) using the R software (v 3.4.3) and the “RVAidememoire” package (Hervé, 2014) (https://cran.r-project.org/web/packages/RVAideMemoire/index.html)(script is available on Zenodo at https://doi.org/10.5281/zenodo.7018260). PLS-DA is an appropriate analysis when the dataset contains fewer groups than explanatory variables, as in the case of the present quantitative dataset. PLS-DA is a supervised technique, meaning that class memberships of the CHCs need to be predefined. Here, we only used the first eight axes produced by the PLS to perform two PLS-DA tests: the first analysis considered the four population samples as four different classes (LCG, GCG, LiPF, GPF, outlined on Fig. 1). The second analysis was based on two classes only, representing the two host types (PF, CG). The number of significant PLS components was determined by cross model validation. We also calculated a numerical value representing the importance of each CHC variable in the projection (VIP), i.e. VIP values larger than 1 are most influential (Hervé, 2014).

## Results

### Hydrocarbon profiles

We detected a complex pattern of 22 hydrocarbons and two non-identified components (I1 and I2) in the cuticle of *I. uriae* ticks. The hydrocarbon mixture was composed of: 14 *n*-alkanes ranging from 17 to 33 carbon atoms in length, 4 monomethyl-alkanes with 17 to 24 carbon atoms and 4 alkenes with 21 to 27 carbon atoms (Table 1).

**Table 1:** List of detected compounds in the cuticle of *I. uriae* adult female ticks, including name and type.

| Cuticular component | Name | Type |
| --- | --- | --- |
| C<sub>17</sub> | *n*-heptadecane | alkane |
| meC<sub>17</sub> | methylheptadecane | methyl-branched alkane |
| C<sub>18</sub> | *n*-octadecane | alkane |
| C<sub>20</sub> | *n*-eicosane | alkane |
| C<sub>21 :1</sub> | heneicosene | alkene |
| C<sub>21</sub> | *n*-heneicosane | alkane |
| C<sub>22 :1</sub> | docosene | alkene |
| C<sub>22</sub> | *n*-docosane | alkane |
| 2meC<sub>22</sub> | 2methyldocosane | methyl-branched alkane |
| C<sub>23</sub> | *n*-tricosane | alkane |
| C<sub>24</sub> | *n*-tetracosane | alkane |
| 9meC<sub>24</sub> | 9methyltetracosane | methyl-branched alkane |
| 2meC<sub>24</sub> | 2methyltetracosane | methyl-branched alkane |
| C<sub>25 :1</sub> | pentacosene | alkene |
| C<sub>25</sub> | *n*-pentacosane | alkane |
| C<sub>26</sub> | *n*-hexacosane | alkane |
| C<sub>27 :1</sub> | heptacosene | alkene |
| C<sub>27</sub> | *n*-heptacosane | alkane |
| C<sub>28</sub> | *n*-octacosane | alkane |
| C<sub>29</sub> | *n*-nonacosane | alkane |
| C<sub>31</sub> | *n*-hentriacontane | alkane |
| C<sub>33</sub> | *n*-tritriacontane | alkane |
| I1 | Non-identified | - |
| I2 | Non-identified | - |

All hydrocarbons were shared by the four tick populations, except 2meC_24_ and C_33_ which were only found in puffin ticks (Fig.2).

**Figure 2:**
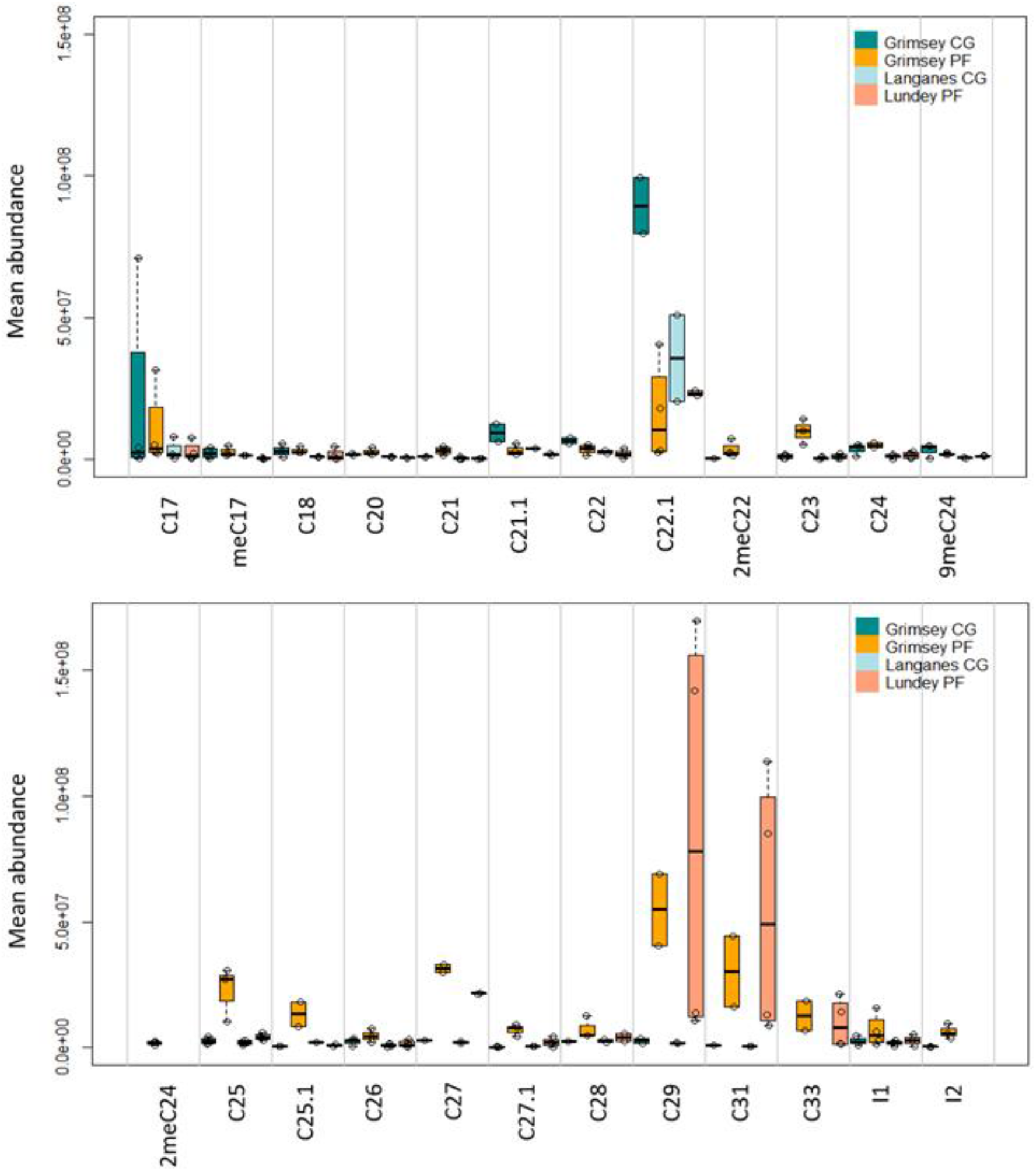
Boxplot representing the mean abundance of each detected cuticular hydrocarbon from *I. uriae* ticks. Compounds ranged from 17 to 33 carbon atoms in length. Mean abundance was calculated based on values from four replicate tick pools. First and third quartiles, median and data points are shown. See Table 1 for CHC abbreviations. PF refers to the Atlantic puffin (*Fratercula arctica*) and CG to the Common guillemot (*Uria aalge*).

A high degree of variation among samples in mean abundance of cuticular components was obvious (Fig.2). Hydrocarbons C_22:1_, *n-*C_29_ and *n-*C_31_ were the most predominant, whereas many hydrocarbons were detected in low quantities, as for example *n-*C_20_, 2meC_22_ and *n-*C_26_.

The abundance pattern of cuticular hydrocarbons varied between ticks of the two host species (Fig.3), but also among the same host in different colony sites. Nevertheless, results of all pairwise profile comparisons were non-significant. No CHC was specific to CG samples, although *n-*C_17_ and C_22:1_ were highly abundant in the GRI colony (Fig.2 and 3). In contrast, long chain cuticular hydrocarbons tended to be present in both PF samples: *n-*C_27_, *n-*C_29_ and *n-*C_31_. CHC components tended to be in lower overall abundance in samples from Langanes (Langanes CG), whereas samples from Lundey Island (Lundey PF) had the highest overall abundance (Fig. 2).

**Figure 3:**
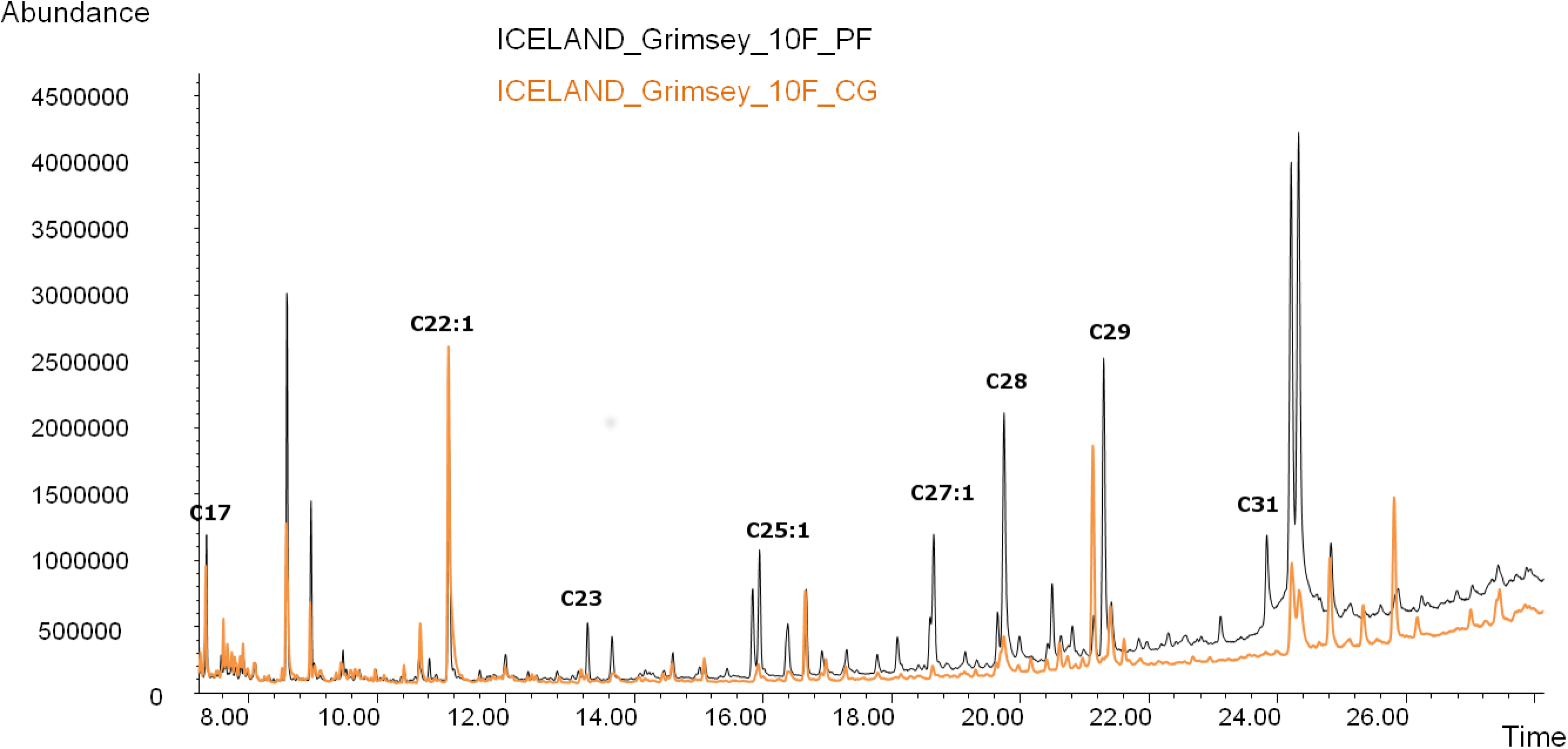
Gas-chromatograms showing the comparison of cuticular hydrocarbon profiles for pools of 10 female ticks from Atlantic puffins (PF: black line) and Common guillemots (CG: orange line) on Grimsey Island, Iceland.

### PLS-DA analyses

#### Population samples

Cross model validation for population samples showed that 50.6% of samples were assigned to the population of origin. PLS-DA analysis showed a discriminating population effect (p=0.008) separating the two host species on first axis (Fig.4). The three first axes explained respectively 11.94, 7.99 and 3.99 % of the total variance among samples.

**Figure 4:**
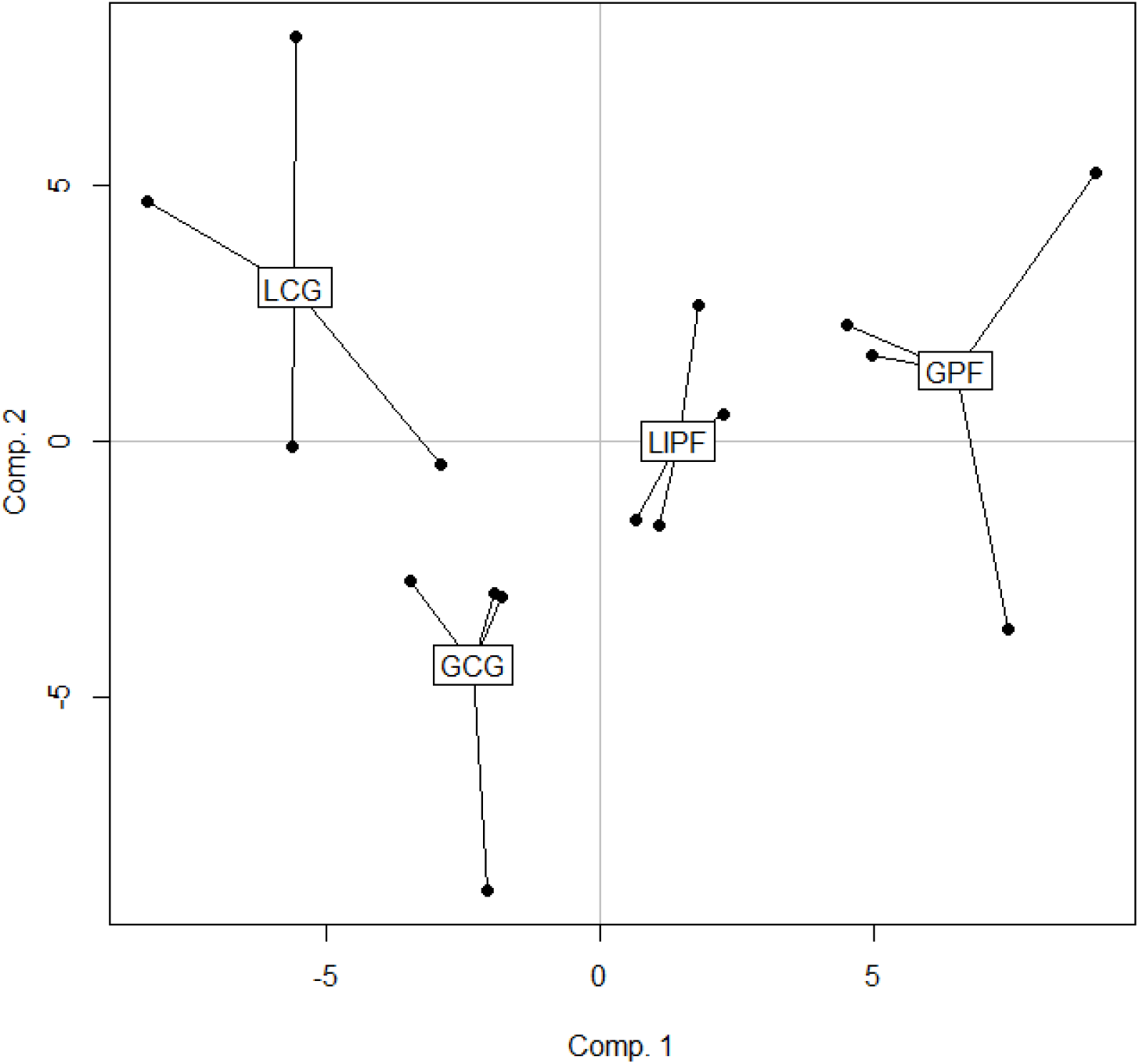
Graphical representation of the four population samples on the two first axes of the PLS-DA analysis. The first and second axes explained respectively 11.94 and 7.99% of the total variance among samples (p=0.008). The acronyms of the sampling locations are as follows: Grimsey - G; Langanes - L; Lundey Island - Li. PF refers to Atlantic puffin (*Fratercula arctica*) and CG to Common guillemot (*Uria aalge*).

Hydrocarbons *n*-C_33_, 2meC_24_ and *n*-C_29_ appeared as the most influential components separating the four population samples (Fig.5).

**Figure 5:**
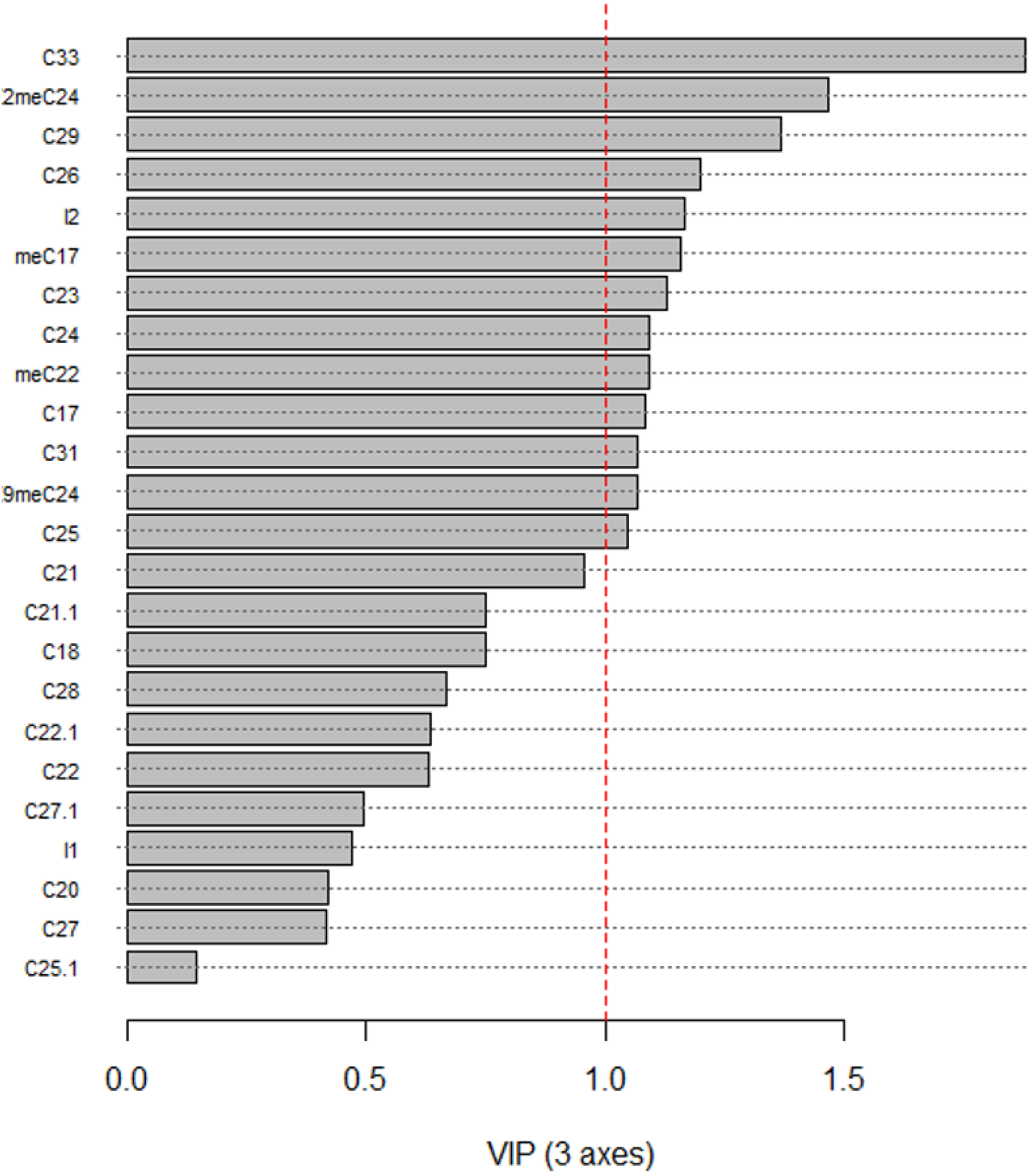
VIP classification showing the influence of CHC in discriminating the four population samples. VIP values larger than 1 (dashed red line) are most influential. See Table 1 for CHC abbreviations.

#### Host types

The cross-validation test for host type revealed that 75% of the tick pools were assigned to the host of origin (PF, CG). However, the PLS-DA analysis revealed no significant difference in CHC profiles between the host types, although a tendency was evident (PLS-DA: p=0.064).

## Discussion

### Cuticle composition and function

Cuticular hydrocarbons, including linear and methyl-branched alkanes and alkenes, are biosynthesized from an acetyl-CoA molecule in specialized secretory cells, the oenocytes which are mainly located in the epidermis of insects (Howard and Blomquist, 2005). They are shuttled through the hemolymph to the epicuticular surface via specialized pore canals penetrating the cuticular layers (Holze et al, 2021). Once fixed on the arthropod cuticle within a complex mixture of alcohols, esters, aldehydes, fatty acids, etc, hydrocarbons help prevent desiccation and serve in chemical communication, constituting essential mediators of insect behavior (Blomquist and Bagnères, 2010). Using GC-MS techniques to analyze extracts from female *I. uriae* ticks, we detected a complex mixture of cuticular hydrocarbons containing linear and monomethyl-alkanes and alkenes ranging from 17 to 33 carbon atoms. The qualitative composition is similar to that reported for many other arthropods, comprising predominantly linear alkanes (*n*-C_23_, *n*-C_25_, *n*-C_27_, *n*-C_29_ and *n*-C_31_) (Howard and Blomquist, 2005; Lockey, 1988). A majority of the detected cuticular hydrocarbons were already reported in other hard ticks: *I. persulcatus* (Tkachev et al., 2000), *Amblyomma variegatum* (Estrada-Peña et al., 1994) and *Rhipicephalus* spp. (Estrada-Peña et al., 1992). Nevertheless, alkenes were only detected once in low quantities in *Rhipicephalus* spp. (Estrada-Peña et al., 1992). Here, we report the presence of alkenes as heneicosene, pentacosene, heptacosene and a high quantity of docosene, particularly in one of the CG samples. Alkenes were demonstrated to act as sex pheromones in the Alfalfa leaf-cutter bee *Megachile rotundata* and the rove beetle *Aleochara curtula* (Paulmier et al., 1999; Peschke and Metzler, 1987). Heneicosene is also described as an aggregation pheromone in *Drosophila* species (Bartelt et al., 1988; Bartelt and Jackson, 1984). Pentacosene is implicated in the mating process of different fly species as a stimulant pheromone (Uebel et al., 1978). The presence of these alkenes in *I. uriae* could corresponds to the biological state of the female ticks when they were collected. However, as mating in this tick species frequently takes place prior to feeding on the host, and in particular when nymphal ticks emerge as flat females (McCoy and Tirard, 2002), we cannot conclude on the mating status of collected females. Indeed, some of the profile variation in our data may be due to the inclusion of females in different reproductive states. Experimental studies will now be necessary to determine the potential role of the observed CHC compounds in mating behavior.

We found also large amount of long-chain hydrocarbons in PF samples (*n*-C_29_, *n-*C_31_ and *n*-C_33_). Mixed with other compounds, *n*-C_31_ was demonstrated to induce copulation in males of the stable fly *Stomoxys calcitrans* (Uebel et al., 1975). Other long chain hydrocarbons, including *n-*C23 to *n-*C_31_, produced in large quantities and acting in combination, were found to serve in colony recognition by bumblebees *Bombus terrestris* (Rottler et al., 2013). Studies have also highlighted the importance of the environment in the acquisition of new hydrocarbons (d’Ettorre et al., 2006; d’Ettorre et al., 2002), acting for example as a recognition signal of their own nest for the social wasp *Polistes metricus* Say (1981) (Singer and Espelie, 1996; Espelie et al., 1990). Moreover, 2meC_24_, which we only detected in PF CHC profiles, has been shown to serve as a contact pheromone in peach twig borers *Anarsia lineatella* (Schlamp, 2005). As puffin burrows are often deep, and densely distributed (Harris and Wanless, 2011), the production of these cuticular hydrocarbons in *I. uriae* may enable ticks to find conspecifics in the host nesting environment. The use of this particular blend of CHCs may also help ticks to find individuals that smell similar, favoring assortative mating (van Zweden and d’Ettorre, 2010). The production of a specific CHC blend may not be necessary for CG ticks because guillemots breed in extremely dense numbers on cliff ledges with no constructed nest; ticks tend to aggregate under and around rocks on the cliff ledge such that finding a mate from the same host type may be easier compared to ticks exploiting puffin hosts. More generally, the presence of complex compounds in the CHC pattern of *I. uriae* suggests that chemical communication may be important in this tick species, potentially enabling ticks to find a suitable mating partner that would enable them to maintain host-adapted gene complexes (Sonenshine and Roe, 2014).

### Alternative hypotheses to explain the origin of CHC patterns

As expected, chemical analyses revealed that each tick population had a distinct CHC profile, but that specific CHCs were also associated with different host types. In particular, *n*-C_33_, 2meC_24_ and *n*-C_29_ were most frequently or exclusively detected in PF populations. The quantitative variability among detected hydrocarbons could be related to different factors.

First, aging and development have been demonstrated to impact cuticular hydrocarbon patterns in different taxa (Desena et al., 1999; Ichinose and Lenoir, 2009). For example, aging favors the production of longer hydrocarbon chains and decreases sexual attractiveness in *Drosophila melanogaster* (Kuo et al., 2012). The high quantities of long-chain hydrocarbons (*n*-C_27_, *n*-C_29_, *n*-C_31_) observed in the puffin tick samples could be a consequence of age variability between host-associated tick samples. Samples used in the present study were collected in the field and the relative age of the specimens could not be determined. However, as tick activity is relatively synchronous with seabird breeding, we did not expect the overall timing in adult female activity to differ in a systematic way between ticks exploiting the different host species, nor among distinct colony locations within Iceland.

Second, climatic conditions can also shape CHC composition. Differences in cuticular hydrocarbon composition were observed in geographically distant tick populations of *Amblyomma variegatum* (Estrada-Peña et al., 1994) and some extreme climatic parameters were shown to be correlated with methyled-alkanes in *Rhipicephalus sanguineus* (Estrada-Peña, 1993). These results suggest a potential link between variation in CHC compounds and adaptation to environmental temperatures. The different types of off-host environments used by *I. uriae* (rock or burrow) may display significant variation in terms of temperature and relative humidity due to differential exposure to climatic factors such as sun, rain and/or snow (Buckley and Buckley, 1980). These selective factors can impact tick survival and could lead to the quantitative variation in cuticular components observed in this study. Howard et al (1978) argued that the effectiveness of CHCs in preventing desiccation depends on the quantity of CHCs and that saturated CHCs are more effective components in protecting against water loss. By positioning temperature and humidity captors during one year in the off-host environment in a heterospecific seabird breeding colony in northern Norway (70°22’ N, 31°10’ E), we observed that the average temperature and relative humidity (HR) ranged respectively from −7.5 to 18°C and from 0 to 110% HR in breeding sites of Common guillemots, and from −7.5 to 10°C and from 80 to 105% HR in those occupied by puffins (data not shown). The micro-habitat used by CG ticks may therefore be more exposed to temperature and humidity variation than the more stable deep burrows used by PF ticks. However, the presence and abundance of saturated CHCs, such as 9meC_24_ and 2meC_24_, were not more frequent in CG samples compared to PF samples. More detailed environmental data from each location and more sampled locations are therefore necessary to more fully evaluate this hypothesis.

Third, diet also appears to be an important factor shaping cuticular hydrocarbon profiles. In the Argentine ant *Linepithema humile*, colonies eating different prey items present particular hydrocarbon profiles that include components coming directly from the prey (Liang and Silverman, 2000). In the same way, Geiselhardt et al., (2012) showed that males of the phytophagous mustard leaf beetle *Phaedon cochleariae* preferred to mate with females reared on the same host plant compared to females from a different host plant, even though they originated from the same laboratory stock population. Studies have also shown that the ectoparasite *Varroa destructor* can mimic its host’s cuticular hydrocarbons, enabling the parasite to escape the hygienic behavior of the host honeybee (Le Conte et al., 2015; Nazzi and Le Conte, 2016). In ticks, it was demonstrated that the cuticular fatty acid profiles of *Rhipicephalus* spp. presented significant differences in abundance according to host use (Shimshoni et al., 2013). In our system, avian erythrocyte membrane lipids might provide a direct pathway for the tick to synthesize cuticular hydrocarbons from pre-existing shorter-chain fatty acids and could result in the acquisition of specific hydrocarbon mixtures. This type of acquisition could explain both the variation among ticks from different host species, and variation among colony locations, if seabird diets shift among locations. Future studies could therefore be expanded to look for correlations between avian erythrocyte membrane lipid composition and tick cuticular signatures by comparing ticks and host blood samples. In addition, looking at patterns in tick cuticular lipids, such as fatty acids and steroids, which have been reported as pheromones in metastriate ticks (Sonenshine, 2004; Sonenshine et al, 1985), would also be of particular interest.

Fourth, CHCs can be altered by infection with different micro-organisms. For example, both the corn borer (*Ostrinia nubilalis*) and the common cockchafer (Melolontha melolontha) show altered CHC profiles when infected by fungi of the genus Beauveria (Lecuona et al, 1991). Moreover, Yoder and Domingus (2003) demonstrated that production of the CHCs *n-*C_20_ and *n-*C_24_ by *Dermacentor variabilis* ticks could protect against predatory behavior by ants. The abundances of *n-*C_20_, *n-*C_21_, *n-*C_23_ and *n-*C_24_ in the puffin tick pools of the present study may represent different infection statuses and/or that puffin burrows provide a less protective environment against predators, such as ants or spiders, than stones in common guillemot colonies.

In general, making inferences on environmental versus host effects will require the examination of ticks from additional sites and host types. However, regardless of the origin of host-associated differences in CHC profiles of *I. uriae*, it is important to now determine is whether these differences reinforce assortative mating patterns, favoring the divergence of sympatric populations and the rapid formation of local host races. Future analyses should focus on the characterization and isolation of the main components of the cuticular mixture from ticks of each seabird host type to test for their biological activity and potential role in tick mating behavior.

## Acknowledgements

We would like to thank Yann Kolbeinsson from Northeast Iceland Nature Research Centre, Iceland for sampling help. We thank “Tiques et Maladies à Tiques” Working Group of the Societé Française d’Ecologie et d’Evolution (SFE2) for stimulating discussions and support.

Version 5 of this preprint has been peer-reviewed and recommended by Peer Community In Zoology (https://doi.org/10.24072/pci.zool.100014).

## Data, scripts and codes availability

Data are available online: https://doi.org/10.5281/zenodo.6497483

## Supplementary material

Supplementary material are available online: https://doi.org/10.5281/zenodo.5889077

